# *C. elegans* Hedgehog-related proteins are tissue- and substructure-specific components of the cuticle and pre-cuticle

**DOI:** 10.1101/2023.12.26.573316

**Authors:** Nicholas D. Serra, Chelsea B. Darwin, Meera V. Sundaram

## Abstract

In *C. elegans*, divergent Hedgehog-related (Hh-r) and Patched-related (PTR) proteins promote numerous processes ranging from epithelial and sense organ development to pathogen responses to cuticle shedding during the molt cycle. Here we show that Hh-r proteins are actual components of the cuticle and pre-cuticle apical extracellular matrices (aECMs) that coat, shape, and protect external epithelia. Different Hh-r proteins stably associate with the aECMs of specific tissues and with specific substructures such as furrows and alae. Hh-r mutations can disrupt matrix structure. These results provide a unifying model for the function of nematode Hh-r proteins and highlight ancient connections between Hh proteins and the extracellular matrix.

## Introduction

*C. elegans* has no ortholog of Hedgehog (Hh) or most of its canonical signaling partners. However, the *C. elegans* genome encodes more than 60 secreted proteins thought to be evolutionarily related to Hh across four families: Warthog (WRT), Groundhog (GRD), Groundhog-like (GRL), and Quahog (QUA) (Bürglin 1996; Bürglin and Kuwabara 2006; Hao *et al*. 2006c; Bürglin 2008a; b). Some of these proteins contain a HOG/Hint domain similar to the autocatalytic C-terminus of Hh. However, none have any sequence similarity to the signaling domain of Hh; instead, they contain nematode-specific cysteine domains for which they are named (WRT, GRD, GRL, or QUA). The PTR protein family, related to the Hh receptor in other systems, has undergone coincident expansion in nematodes, suggesting that Hh-r and PTR proteins act together in some way (Zugasti *et al*. 2005; Bürglin and Kuwabara 2006; Zhang and Beachy 2023). There is evidence for potential signaling roles of a few Hh-r and PTR proteins, though the relevant downstream pathways remain unclear (Lin and Wang 2017; Kume *et al*. 2019; Templeman *et al*. 2020; Chiyoda *et al*. 2021; Wang *et al*. 2023; Emans *et al*. 2023). PTR proteins can also affect Hh-r endocytosis (Zugasti *et al*. 2005; Chiyoda *et al*. 2021). Hh-r and/or PTR proteins have been tied to numerous functions in development and physiology, but the most abundant evidence points to roles in the organization of apical extracellular matrices (aECMs) (Bürglin and Kuwabara 2006; Cohen and Sundaram 2020).

*C. elegans* external epithelia are covered by a collagenous cuticle aECM that is shed and replaced at each larval molt (Page and Johnstone 2007). Each cuticle is preceded by a transient pre-cuticle aECM that helps pattern the new cuticle and allows proper release of the old cuticle (Cohen and Sundaram 2020). Like cuticle and pre-cuticle genes, many Hh-r genes are expressed in external epithelia and show an oscillatory pattern of gene expression during the molt cycle (Hao *et al*. 2006c; Hendriks *et al*. 2014; Meeuse *et al*. 2020). QUA-1 (the only member of the QUA family) binds cuticle and loss or knockdown of *qua-1* and several *wrt* or *ptr* genes causes cuticle or molting defects (Zugasti *et al*. 2005; Hao *et al*. 2006a; b; Baker *et al*. 2021). Recently, we reported that loss of *ptr-4* disrupts organization of the pre-cuticle aECM, which can explain the *ptr-4* mutant phenotypes (Cohen *et al*. 2021). Fung *et al*. (2023) also reported that GRL-18 localizes transiently to developing cuticle pores over sensory glia, suggesting GRL-18 could be a glial socket pre-cuticle factor. Finally, Chiyoda *et al*. (2021) reported a transient embryonic expression pattern for GRL-7 that we found reminiscent of known pre-cuticle components (Vuong-Brender *et al*. 2017; Birnbaum *et al*. 2023). Here we demonstrate that these and other Hh-r proteins are stable, tissue- and substructure-specific components of the cuticle or pre-cuticle.

## Methods

### *C. elegans* Maintenance and Strains

Animals were grown at 20° C under standard conditions (Brenner 1974). Strains used are shown in Table S1. GRL-2 and WRT-10 fusions were made by Suny Biotech (Fuzhou, China) using CRISPR/Cas9 genome editing. GRL-18 and GRL-7 fusions were kind gifts of Maxwell Heiman (Harvard U.) and Masamitsu Fukuyama (U. Tokyo).

### Confocal microscopy

Confocal images were captured with a Leica TCS DMi8 confocal microscope and Leica Las X Software. Worms were immobilized with 10 mM levamisole in M9 buffer and mounted on 2% agarose pads supplemented with 2.5% sodium azide. L4 animals were staged based on vulva morphology (Mok *et al*. 2015). For each strain, at least n=10 animals were imaged at each stage shown in the figures and in 24-hour adults.

### Fluorescence Recovery After Photobleaching (FRAP)

FRAP experiments were conducted using a Leica DMi8 laser scanning confocal microscope and the Leica Las X Software FRAP module. Pre and post bleach laser powers ranged from 1-2% depending on protein target. A 3µm x 3µm ROI was designated for each experiment. 20 pre-bleach frames were captured at 1 frame/ 0.4s, followed by 20 to 30 bleaching frames at 100% laser power power at 1 frame/ 0.4s depending on target. Recovery timecourse was 3 minutes with image capture at a rate of 1 frame/ 2 seconds for 90 recovery frames total. Replicate FRAP data was analyzed in Graphpad Prism software by fitting one-phase association curves to the data reliant on the following formula Y = Y0 + (Plateau-Y0)*[1–e(2Kx)]. Mobile fraction was calculated by subtracting the Y0 value from Plateau.

### Alae observation

The alae in late L4 and young adult worms were observed by differential interference contrast (DIC) microscopy and by epifluorescence after staining with lipophilic DiI (Biotium) as in (Schultz and Gumienny 2012). Briefly, young adult worms were washed and pelleted from plates with M9 buffer and incubated in 30 µg/ml DiI shaking at 20 °C for 3 hours. After washing again with M9, worms were allowed to recover on OP50 seeded NGM plates for at least 30 minutes before mounting and imaging. Worms were observed with a Zeiss Axioskop (Carl Zeiss Microscopy) with an attached Leica DFC360 FX camera. Images were acquired using the software package Qcapture (Qimaging). Image cropping, scaling, and adjustment was handled in ImageJ.

## Results and Discussion

To test if Hh-r proteins are found within the aECM, we systematically evaluated the localization patterns of 4 Hh-r proteins (GRL-2, GRL-7, GRL-18, and WRT-10) in larvae and adults using endogenously tagged fluorescent reporters. GRL-2 and GRL-18 fusions permanently labelled the cuticle of specific tube cells (Figures 1 and 2), while GRL-7 and WRT-10 fusions transiently labelled distinct substructures within epidermal and seam pre-cuticles (Figures 3 and 4). WRT-10 also labelled the pre-cuticles of specific interfacial tubes (Figure 4). The tissue distribution of each Hh-r protein was consistent with each gene’s reported transcriptional pattern based on transgenic reporters and/or single cell RNA sequencing (Soulavie *et al*. 2018; Packer *et al*. 2019; Fung *et al*. 2020, 2023). We conclude that Hh-r proteins label aECMs of those cells in which they are expressed.

**Figure 1.**
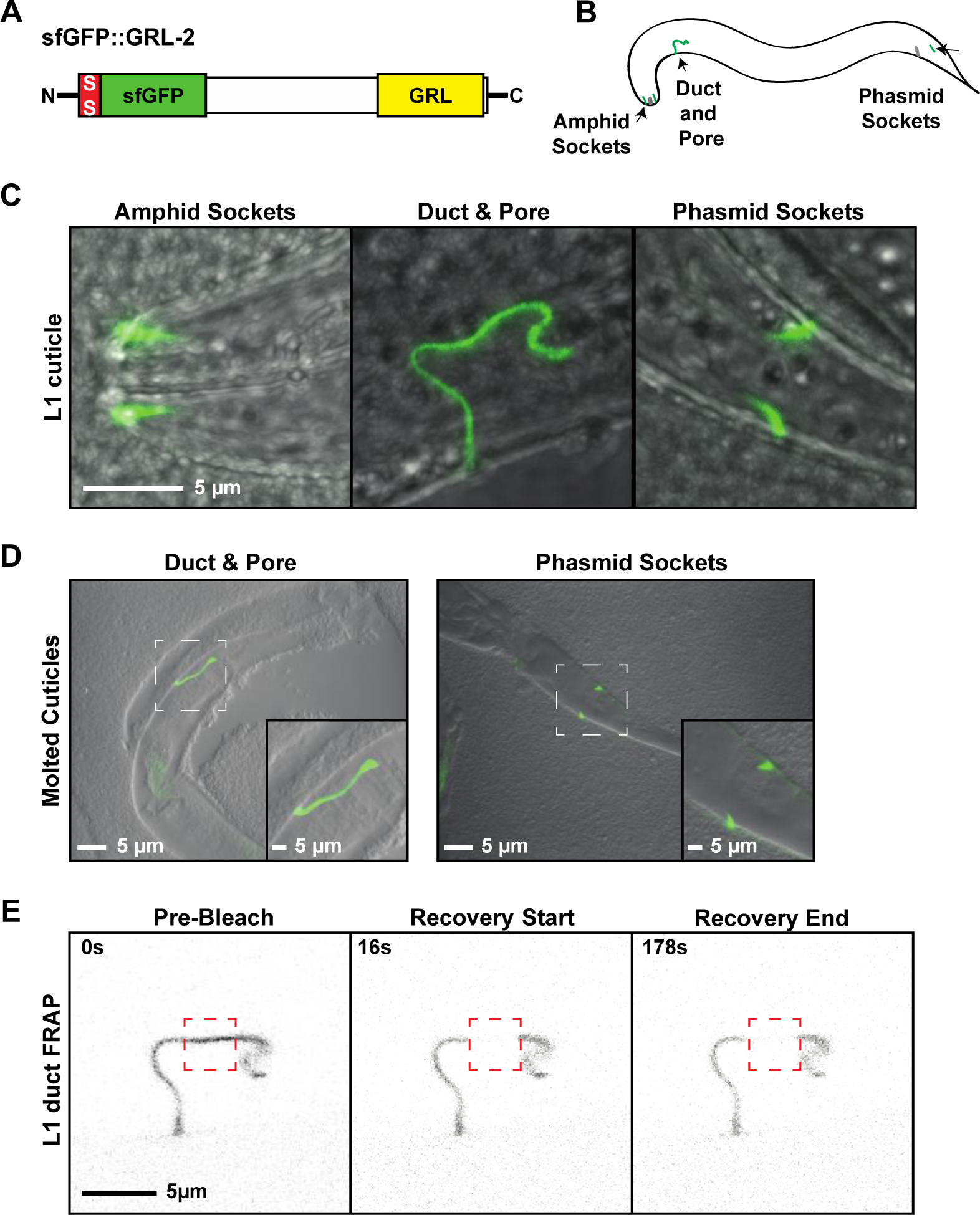
GRL-2 is a tube specific cuticle component. A. SfGFP::GRL-2 schematic. SfGFP was inserted immediately following the signal sequence (SS). B. Cartoon summary of expression pattern. C. SfGFP::GRL-2 labelled the cuticle of the excretory duct and pore tubes and the amphid and phasmid socket glia at all larval stages and in adults. D. It could also be detected in shed cuticles after a molt. E. SfGFP::GRL-2 showed no recovery after FRAP (n=12). See Table 1 for analysis.

**Figure 2.**
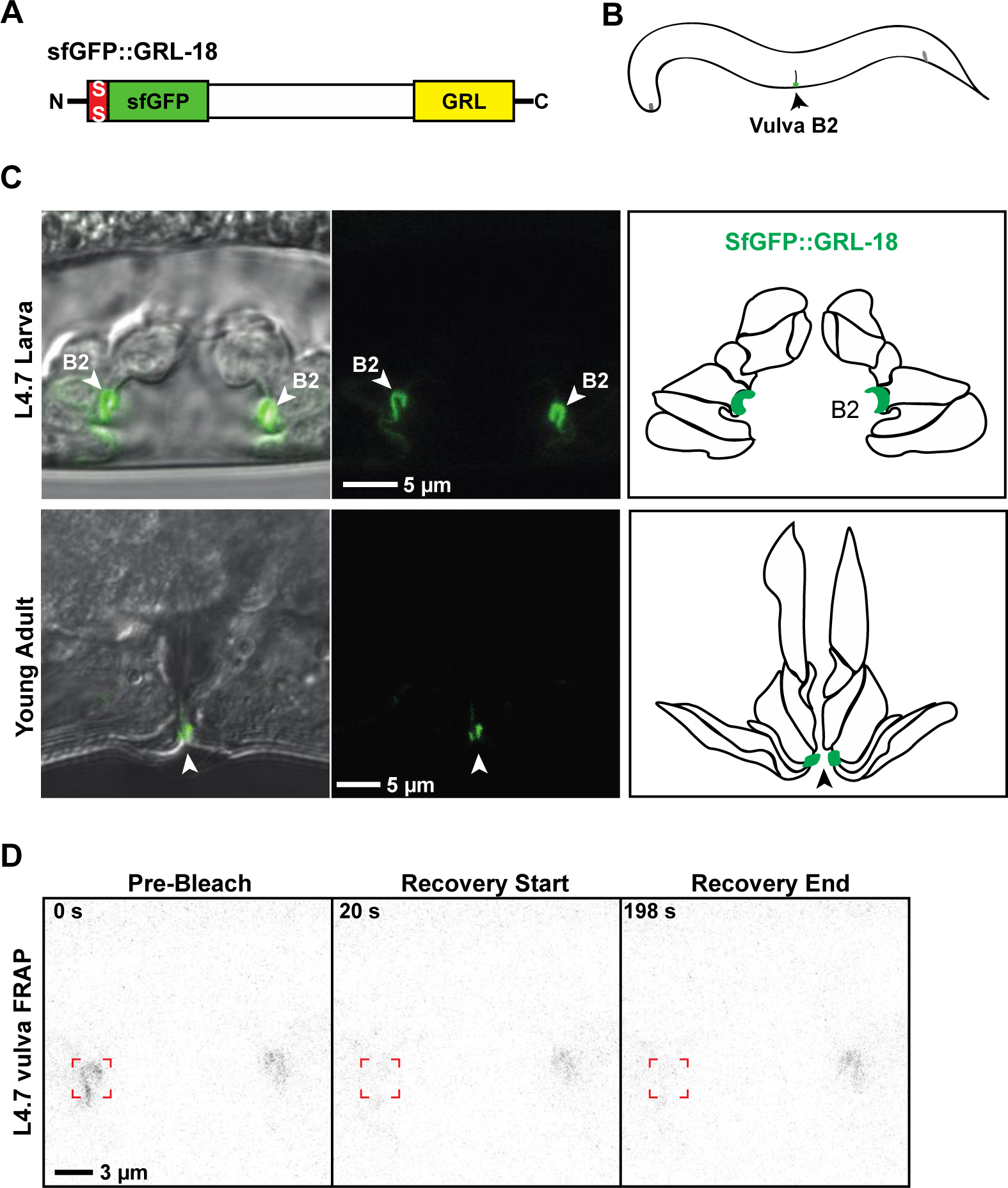
GRL-18 is a cell-specific cuticle component in the vulva. A. SfGFP::GRL-18 schematic and B. Cartoon summary of expression pattern. C. In addition to labelling specific socket glial pores transiently in larvae as previously reported (Fung *et al*. 2023), SfGFP::GRL-18 labelled the cuticle of the vulB2 cell in the hermaphrodite vulva, beginning at the mid-L4 stage and continuing in the adult. Cartoons indicate the seven vulva cell types and the position of vulB2 cells at each stage, based on (Cohen *et al*. 2020). D. SfGFP:;GRL-18 showed no recovery after FRAP (n=15). See Table 1 for analysis.

**Figure 3.**
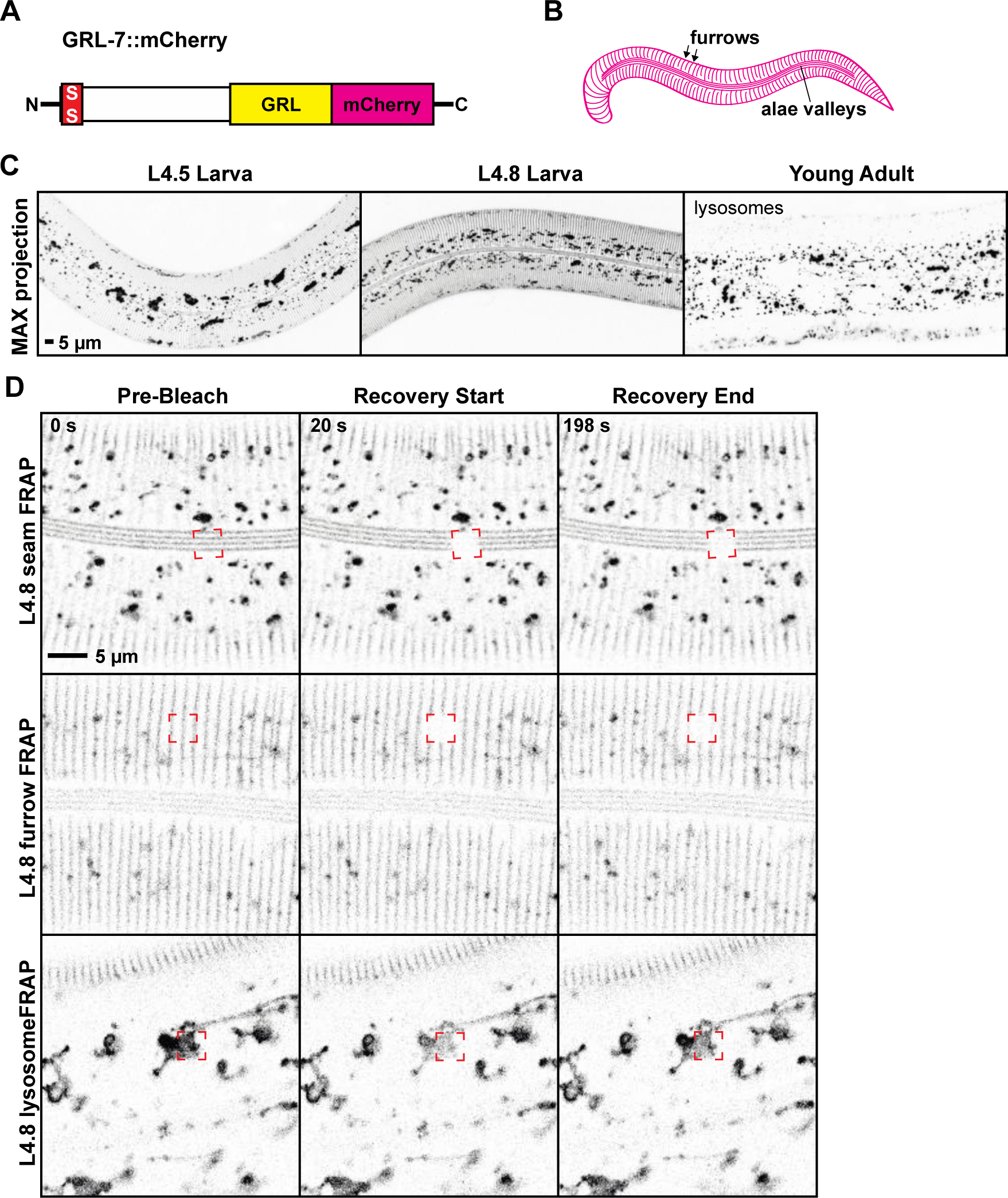
GRL-7 is a furrow-specific component of pre-cuticle. A. GRL-7::mCherry schematic and B. Cartoon summary of expression pattern. C. GRL-7::mCherry transiently labelled developing furrow structures in the epidermal aECM and valley regions between developing alae ridges over the seam. GRL-7::mCherry labelled these aECM structures from the L4.5 to L4.9 stages and then was endocytosed and cleared before the molt. In adults, it was observed only in internal lysosome-like structures, where mCherry fusions often persist (Clancy *et al*. 2023; Birnbaum *et al*. 2023). This pattern of expression and endocytosis is similar to that reported for other pre-cuticle factors (Birnbaum *et al*. 2023) and for GRL-7 in embryos (Chiyoda *et al*. 2021). D. In both the seam and epidermal aECM, GRL-7::mCherry showed no recovery after FRAP (n=11 each). However, lysosomal signal recovered rapidly (n=6). See Table 1 for analysis.

**Figure 4.**
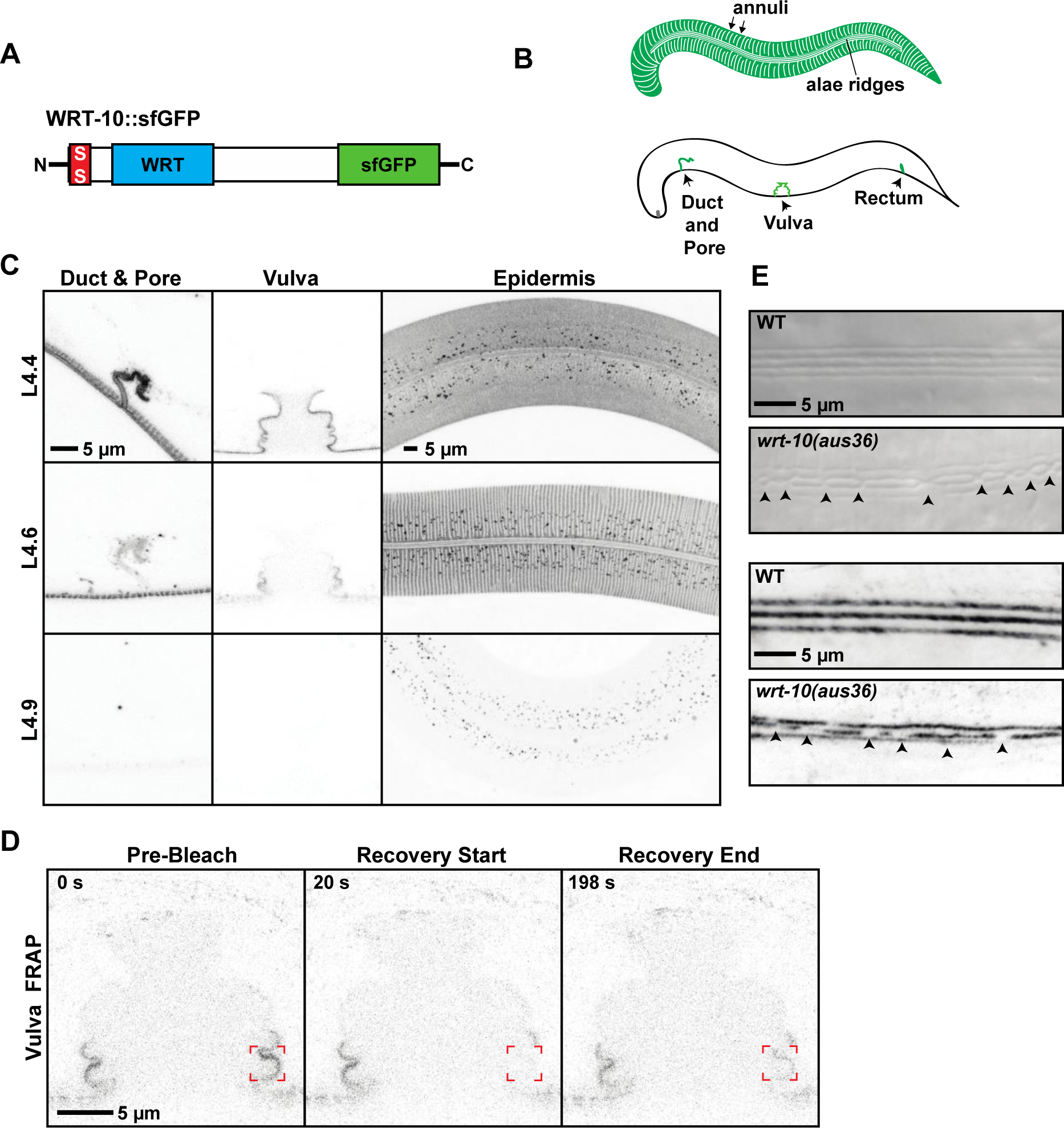
WRT-10 is a broadly expressed component of pre-cuticle. A. WRT-10::SfGFP schematic and B. Cartoon summaries of expression pattern. The epidermal and tube expression are coincident but are shown separately for clarity. C. WRT-10::SfGFP transiently labelled the excretory duct and pore, rectum (not shown), and vulva tubes in the intermolt periods, along with developing annuli structures in the epidermal aECM and developing alae ridges over the seam - a pattern complementary to that of GRL-7 (see Fig. 3). Like GRL-7, WRT-10::SfGFP was endocytosed and cleared before the molt. It was not detectable in adults. D. In the vulva, WRT-10::SfGFP showed limited recovery after FRAP. See Table 1 for analysis. E) *wrt-10(aus36)* adults have fragmented alae ridges (n=17/17), shown here with DIC (top) and after DiI staining, which specifically labels ridges and not valleys (Schultz and Gumienny 2012). Similar results were seen with *wrt-10(aus37)* (n=15/16).

**Table 1.** FRAP analyses.

| **FRAP Experiment** | **n** | **Y0** | **Plateau** | **Half-time** | **Mobile Fraction** |
| --- | --- | --- | --- | --- | --- |
| sfGFP::GRL-2 in Duct Lumen | 12 | 0.1492 | 0.1977 | 8.839 | 0.0485 |
| sfGFP::GRL-18 in Vulva Lumen | 15 | 0.5214 | 0.5436 | 56.89 | 0.0222 |
| GRL-7::mCherry Alae Valleys | 11 | 0.2057 | 0.274 | 15.95 | 0.0683 |
| GRL-7::mCherry Furrows | 11 | 0.2747 | 0.3418 | 40.58 | 0.0671 |
| GRL-7::mCherry Lysosomes | 6 | 0.1023 | 0.5242 | 14.3 | 0.4219 |
| WRT-10::sfGFP Vulva Lumen | 12 | 0.2754 | 0.542 | 30.64 | 0.2666 |

Values were calculated from fitted one-phase association curves (see Methods).

To test if Hh-r proteins are stably associated or labile within the aECM, we performed fluorescence recovery after photobleaching (FRAP). GRL-2, GRL-18, and GRL-7 all showed essentially no recovery in these experiments (Figures 1E, 2D, 3D, 4D and Table 1). In contrast, lysosomal GRL-7::mCherry recovered rapidly (Figure 3E and Table 1). WRT-10 showed limited recovery (Figure 4D), with a small mobile fraction similar to that previously reported for other pre-cuticle components such as the Zona Pellucida (ZP) protein LET-653 (Gill *et al*. 2016; Forman-Rubinsky *et al*. 2017). Together with the presence of SfGFP::GRL-2 in shed cuticles (Figure 1D), these data are consistent with Hh-r proteins being stably incorporated into aECMs.

To assess the possibility that Hh-r proteins play structural roles in aECM organization, we examined *grl-2, grl-18, grl-7* and *wrt-10* deletion mutants for matrix-related defects. All mutants were viable (Table S2) and most appeared morphologically normal. However, two independent *wrt-10* mutants had fragmented cuticle alae ridges (Figure 4E), a phenotype similar to that seen in multiple other pre-cuticle mutants (Forman-Rubinsky *et al*. 2017; Katz *et al*. 2022). We conclude that Hh-r proteins can play structural roles in aECM organization, although their mutant phenotypes are generally milder than those caused by loss of PTR proteins.

Structural roles of Hh-r proteins within cuticle and pre-cuticle aECM could explain many of their reported functions in tissue shaping, molting, and pathogen responses (Hao *et al*. 2006a; b; Lin and Wang 2017; Baker *et al*. 2021; Zárate-Potes *et al*. 2022). Cuticle damage could also release Hh-r components for signaling to activate stress and immune pathways (Dodd *et al*. 2018; Martineau *et al*. 2021; Wang *et al*. 2023). Furthermore, aECM tethering could help present Hh-r signals at a particular time and place. In this regard, the precise timing at which different pre-cuticle factors are expressed, assembled, and disassembled from matrix (Hendriks *et al*. 2014; Meeuse *et al*. 2020; Cohen and Sundaram 2020; Birnbaum *et al*. 2023) makes pre-cuticle Hh-r proteins ideally positioned to serve as signals of developmental progression to help coordinate the timing of other events during the molt cycle. The ability of *grl-7* and *grd-1* mutants to suppress the failed L1 diapause or developmental timing defects of other mutants (Kume *et al*. 2019; Chiyoda *et al*. 2021; Emans *et al*. 2023) might be explained by such a model. Finally, the presence of specific Hh-r proteins in the cuticle of tube orifices such as the vulva also raises the possibility of inter-animal signaling to mediate social interactions (Weng *et al*. 2023); however, we did not find any obvious mating deficiency in *grl-18* mutant hermaphrodites (Table S3).

Canonical Hh proteins bind matrix proteoglycans and have intriguing similarities to matrix proteases (Roelink 2018; Gude *et al*. 2023; Zhang and Beachy 2023). In mycobacteria, Patched-related RND transporters transport glycolipids and other cargo to form the outer cell envelope, which is a type of aECM (Nikaido 2018; Dulberger *et al*. 2020). Therefore, it appears that interactions with the ECM are an ancient feature of the Hh-r and PTR protein families.

Although *C. elegans* Hh-r proteins are highly divergent from Hh and other types of non-nematode Hh-r proteins (Bürglin 2008a), their expansion suggests broad and important roles in nematode biology. The realization that Hh-r proteins are embedded within cell-specific and temporally dynamic aECMs provides a foundation for better understanding those roles.

## Acknowledgements

We thank Maxwell Heiman and Masamitsu Fukuyama for strains, Wendy Fung and Maxwell Heiman for stimulating discussions, Mara Cowen for assistance with FRAP, and Wormbase for facilitating data retrieval. Some strains were obtained from the Caenorhabditis Genetics Center, which is supported by the NIH Office of Research Infrastructure Programs P40 OD10440. This work was funded by NIH R35GM136315 to M.V.S.

## Supplemental Materials

**Table S1:** Strains used.

| Strain | Genotype | Description and Source | Reference |
| --- | --- | --- | --- |
| **Fluorescent fusions** |  |  |  |
| CHB4638 | *grl-18(syb6299 [SfGFP::GRL-18]) V* | Endogenous fusion, obtained from Max Heiman | (Fung *et al.* 2023) |
| PHX7022 | *wrt-10(syb7022 [SfGFP::WRT-10]) II* | Endogenous fusion, made by Suny Biotech | This work |
| PHX7064 | *grl-2(syb7064 [GRL-2::SfGFP]) V* | Endogenous fusion, made by Suny Biotech | This work |
| UP4276 | *noah-1(mc68 [NOAH-1::mCherry]) I; grl-18(syb6299 [SfGFP::GRL-18]) V* | Made by crossing CHB4638 with ML2482 | This work |
| YB4668 | *grl-7(td261 [GRL-7::mCherry::3xFLAG]) V* | Endogenous fusion, obtained from Masamitsu Fukuyama | (Chiyoda *et al.* 2021) |
| **Mutants** |  |  |  |
| CHB4546 | *grl-18(hmn341) V* | 960 bp deletion, obtained from Max Heiman | (Fung *et al.* 2023) |
| HRN666 | *wrt-10(aus36) II* | 5 bp deletion/frameshift, obtained from CGC | (Sherry *et al.* 2019) |
| HRN667 | *wrt-10(aus37) II* | 2 bp deletion/frameshift, obtained from CGC | (Sherry *et al.* 2019) |
| RB1999 | *grl-7(ok2644) V* | 424 bp deletion, obtained from CGC | (C. elegans Deletion Mutant Consortium 2012) |
| RB5002 | *grl-2(ok5804) V* | G>A causing premature stop codon, obtained from CGC | (C. elegans Deletion Mutant Consortium 2012) |

**Table S2:** Hh-r mutants are viable.

| Strain | Genotype | Live Adults/Total | % Live |
| --- | --- | --- | --- |
| N2 | *wild-type* | 179/179 | 100 |
| CHB4546 | *grl-18(hmn341)* | 178/178 | 100 |
| N2 | *wild-type* | 157/160 | 98 |
| RB1999 | *grl-7(ok2644)* | 149/150 | 99 |
| N2 | *wild-type* | 207/209 | 99 |
| HRN666 | *wrt-10(aus36)* | 201/201 | 100 |

Viability was assessed by allowing 4 or 5 adults to lay eggs for several hours, and then comparing the number of eggs laid to the number of surviving adult progeny 4 days later. No significant differences were observed between mutants and wild-type controls (Fisher's Exact Test).

**Table S3.** *grl-18* mutant hermaphrodites show normal mating efficiency.

| Strains Crossed | # Male Progeny | # Herm Progeny | % Male Progeny |
| --- | --- | --- | --- |
| N2 m X N2 h | 169 | 304 | 36 |
| N2 m X N2 h | 201 | 269 | 43 |
| N2 m X N2 h | 155 | 340 | 31 |
| N2 m X CHB4546 h | 196 | 291 | 40 |
| N2 m X CHB4546 h | 205 | 277 | 43 |
| N2 m X CHB4546 h | 156 | 227 | 41 |

Mating efficiency was assessed by pairing 8 N2 males (m) with 3 hermaphrodites (h, either CHB4546 *grl-18(hmn341)* or N2 controls) for 24 hours, with 3 replicate experiments per genotype. *grl-18* mutant hermaphrodites gave similar numbers of male cross progeny to wild-type controls (ns, Mann-Whitney U Test).

## Notes

### Competing Interest Statement

The authors have declared no competing interest.

